# RETA: An R package for whole exome and targeted region sequencing data analysis

**DOI:** 10.1101/121384

**Authors:** Mengbiao Guo, Jing Yang, Yu lung Lau, Wanling Yang

## Abstract

Whole exome and targeted sequencing have been playing a major role in diagnoses of Mendelian diseases, but analysis of these data involves using many complicated tools and comprehensive understanding of the analysis results is difficult.Here, we report RETA, an R package to provide a one-stop analysis of these data and a comprehensive, interactive and easy-to-understand report with many advanced visualization features. It facilitates clinicians and scientists alike to better analyze and interpret this type of sequencing data for disease diagnoses.

**Availability and implementation:** https://github.com/reta-s/reta/releases

## 1 Introduction

Applying next generation sequencing (NGS) on whole exome or targeted gene panels is becoming a powerful tool for molecular diagnoses of Mendelian diseases. Several tools have been developed to assist data analysis like variant calling, annotation, and filtering or prioritizing of the variants, including KGGseq (Li *et al*., 2012), PriVar (Zhang *et al*., 2013), exomeSuite (Maranhao *et al*., 2014) and GeneCOST (Ozer *et al*., 2015). However, an integrative one-stop tool to facilitate various analyses of these data and provide a comprehensive and interactive report for easy understanding of the results is still lacking.

Here we present an R package, RETA, to meet this demand in a user-friendly and easy-to-understand fashion. The important features of RETA include various in-depth quality control measures, integrative coverage examination and visualization, detection of runs-of-homozygosity and interactive, straightforward analysis results presentation. Inheritance mode consideration allows in-depth processing of the data for variants prioritization for causality analysis.

## 2 Description

### 2.1 Modules

RETA consists of six modules.

- **General QC**. It will provide (1) general summary of the targeted regions, e.g. total number of regions and base pairs according to the selected platform; (2) summary of sequencing data including number of reads, average GC ratio and read length.
- **In-depth QC**. (1) zygosity check for both X chromosome and autosome variants for checking gender misdesignation, sample contamination or consanguinity; (2) identity-by-descent (IBD) analysis for family relationship check or marriage consanguinity; (3) distribution of depth of base pairs in the target regions; (4) summary statistics table of variants including Ti/Tv ratio (transition over transversion), distribution of variant regions and genotyping quality.
- **Candidate gene QC**. This module analyzes the detailed coverage of each candidate gene from a list of user choices and reports the rare variants within each gene. Tables here includes (1) summary of candidate genes containing regions of low coverage (<5X by default), with each gene name hyperlinked to a detailed coverage figure in the last section of the report; (2) an interactive table showing the low coverage regions in all the candidate genes. Similarly, (3) a table showing regions of low mapping quality in candidate genes; (3) rare variants in candidate genes, excluding variants with minor allele frequency (MAF) > 1%, low quality or coverage.
- **Structural variants analysis**. We included runs-of-homozygosity (ROH) regions here because large ROH usually indicates large heterozygous deletions, consanguinity or uniparental disomy. Tables in this section include: (1) ROH regions identified, and homozygous variants found in them. (2) copy number variants (CNVs, large duplications and deletions) called and genes within them. We included some preprocessed exome data for each platform to assist CNV calling, mainly serving as a valid background comparison especially when users have very few samples as input. Each CNV region has a detailed figure in the last section of the report.
- **Variants prioritization based on inheritance modes.** Here we perform variant filtering and prioritization based on assumed inheritance modes, including autosomal recessive (including homozygous or compound heterozygous), autosomal dominant, de novo mutation, and X-linked inheritance. Candidate variants are shown in each subsection based on the corresponding inheritance mode.
- **Detailed figures for CNV and coverage analysis**. This section contains all the figures mentioned in the CNV section and coverage analysis section for candidate genes. For coverage plots, each problematic gene has two figures associated with it, one for the whole gene for an overview of coverage, and the other zoomed-in figure for the regions of low coverage highlighted by vertical rectangles (Figure 1).

**Fig. 1.**
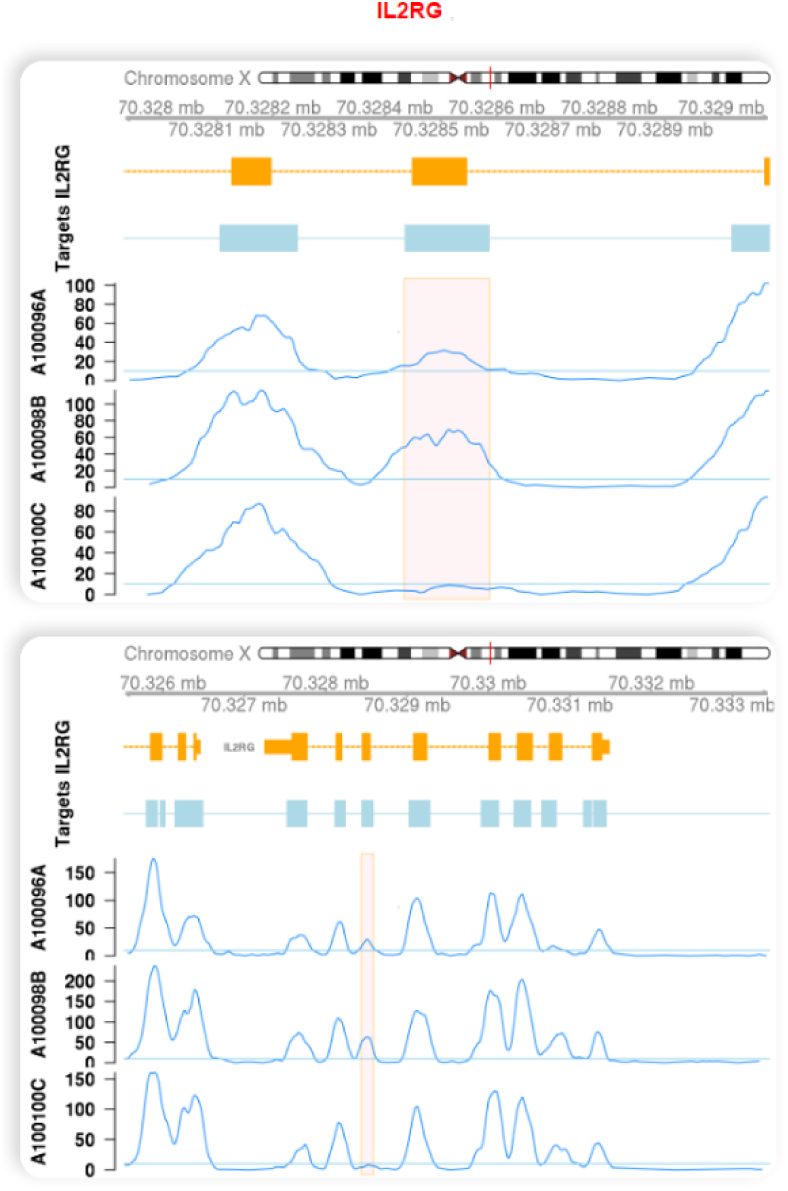
Detailed coverage example for IL2RG gene. (**Top**) Zoomed-in plot of regions of low coverage only. Tracks from top to bottom: chromosome ideogram, genomic coordinates, genes, target regions, coverage for each sample (three samples here, y-axis stands for depth). Highlighted orange rectangles indicate the target regions with low coverage. (**Bottom**) Coverage plot for the whole gene.

### 2.2 Features

As far as we know, this is the first software performing comprehensive quality control and coverage evaluation for candidate gene panels, and providing interactive and integrative report for exome or targeted region sequencing data, with the following features:

- Minimal software requirement: no other tools are needed after installing RETA’s dependent R packages.
- Integrative coverage evaluation and visualization: this points to possible reasons for negative detection and thus new directions to work on, especially when one failed to identify promising candidate variants.
- In-depth quality control: for example, IBD analysis for family member relationship check and variant heterozygosity analysis for sample contamination or gender misdesignation.
- ROH analysis: It allows identifying probable recessive variants from consanguineous family and cases of large heterozygous deletions or uniparental disomy.
- Built-in candidate gene panels and platform designs: users can simply choose the candidate gene panel according to their studied disease and select the platform corresponding to the targeted or exome sequencing platforms used.
- Interactive and straightforward analysis report: well organized results make understanding of the results straightforward.

## 3 Example

Documentation and an analysis report in HTML format for demonstration were included in the supplementary materials.

### Funding

MG is partially supported by the Edward & Yolanda Wong Scholarship. WY and YLL thank support from Research Grants Council of Hong Kong (GRF 17125114, HKU783813M, HMRF 01120846).

*Conflict of Interest*: none declared.

